# Natively Oxidized Amino Acid Residues in the Spinach Cytochrome *b_6_f* Complex

**DOI:** 10.1101/227835

**Authors:** Ryan M. Taylor, Larry Sallans, Laurie K. Frankel, Terry M. Bricker

**Affiliations:** Department of Biological Sciences, Biochemistry and Molecular Biology Section, Louisiana State University, Baton Rouge, LA 70803; The Rieveschl Laboratories for Mass Spectrometry, Department of Chemistry, University of Cincinnati, Cincinnati, OH 45221

**Keywords:** cytochrome *b_6_f* complex, mass spectrometry, Reactive Oxygen Species, spinach

## Abstract

The cytochrome *b_6_f* complex of oxygenic photosynthesis produces substantial levels of reactive oxygen species (ROS). It has been observed that the ROS production rate by *b_6_f* is 10-20 fold higher than that observed for the analogous respiratory cytochrome *bc*_1_ complex. The types of ROS produced (O_2_^•−^,^1^O_2_, and, possibly, H_2_O_2_) and the site(s) of ROS production within the *b_6_f* complex has been the subject of some debate. Proposed sources of ROS have include the heme *bp*, PQ_p_^•−^ (possible sources for O_2_^•−^), the Rieske iron-sulfur cluster (possible source of O_2_^•−^ and/or H_2_O_2_), Chl *a* (possible source of ^1^O_2_) and heme *C_n_* (possible source of O_2_^•−^ and/or H2O2). Our working hypothesis is that amino acid residues proximal to the ROS production sites will be more susceptible to oxidative modification than distant residues. In the current study, we have identified natively oxidized amino acid residues in the subunits of the spinach cytochrome *b_6_f* complex. The oxidized residues were identified by tandem mass spectrometry using the MassMatrix Program. Our results indicate that numerous residues, principally localized near *p*-side cofactors and Chl *a*, were oxidatively modified. We hypothesize that these sites are sources for ROS generation in the spinach cytochrome *b_6_f* complex.

## Introduction

The cytochrome *b_6_f* complex acts as a plastoquinol-plastocyanin (cytochrome C553 in cyanobacteria) oxidoreductase and is similar to cytochrome *bc*_1_ complexes present in heterotrophic organisms. Moderate resolution crystal structures (≈ 3 Å) are available for *b_6_f* complexes of both thermophilic cyanobacteria (*Mastigocladus* (Kurisu et al. 2003) and *Nostoc* (Baniulis et al. 2009)) and a mesophilic green alga (*Chlamydomonas* (Stroebel et al. 2003)). Recently, a 2.5 Å structure of the *Nostoc* protein has been presented (Hasan and Cramer 2014). This higher resolution structure allowed the identification of numerous lipids and intra-protein water molecules that were not identifiable in the earlier structures. The *b_6_f* complex is a symmetric dimer with a molecular mass of 220 kDa containing, within each monomer, 8 subunits: Cyt *f* (PetA), Cyt *b_6_* (PetB), Rieske iron-sulfur protein (PetC), subunit IV (PetD), and four smaller subunits (PetG, PetL, PetM, and PetN). These proteins are associated with 7 prosthetic groups: 2 *c*-type hemes, 2 *b*-type hemes, 1 Fe_2_S_2_ cluster, 1 Chl *a*, and 1 β-carotene. Additionally, a plastoquinol-binding site is present on the *p*-side of the complex and a plastoquinone-binding site is present on the *n*-side of the complex. In cytochrome *bc*_1_ complexes, linear electron transport occurs via a modified Q-cycle mechanism (Crofts et al. 2003). However, it is unclear if this is the case for the cytochrome *b_6_f* complex. The presence of the novel heme *c_n_* and the observation that the complex can participate in cyclic electron transport, accepting electrons from reduced ferredoxin possibly via ferredoxin NADP+ oxidoreductase (which appears to be a subunit of the *in vivo* complex (Zhang et al. 2001)), both argue against a classical modified Q-cycle mechanism. The functions of the Chl *a* and the β-carotene are unclear and these have been hypothesized to play a structural role in *b_6_f* assembly or are possibly required for complex stability (Yan et al. 2008).

In thylakoid membranes, reactive oxygen species (ROS) are produced at a number of sites within the linear electron transport chain including PS II, the *b_6_f* complex and PS I. ROS are formed by the excitation of dioxygen (singlet oxygen, ^1^O_2_), the partial reduction of dioxygen (O_2_^•−^, O_2_^2−^, H_2_O_2_, and ^·^OH), and the partial oxidation of water (H_2_O_2_ and ^·^OH). These are unavoidable byproducts of oxygenic photosynthesis. ROS can oxidatively damage proteins, lipids and nucleic acids (Das and Roychoudhury 2014) and, consequently, places limits on photosynthetic productivity (estimated to be at least 10% based on PS II photoinhibition, alone (Long et al. 1994)). It should also be recognized, however, that ROS also serve as signal molecules which can modulate a variety of cellular processes including stress acclimatization, differentiation and development, programmed cell death and pathogen defense (Mittler 2016).

While ROS generation by PS II has been extensively examined (Kim and Jung 1992; Pospíšil 2009, 2016; Kale et al. 2017), relatively few studies have been performed on the *b_6_f* complex. A number of cofactors within the complex have, however, been proposed as sites of ROS production. Cytochrome *bc_1_*y-type complexes, in general, produce O_2_^•−^ (Lanciano et al. 2013), and production of O_2_^•−^ by the *b_6_f* complex has been observed by EPR spin-trapping spectroscopy (Sang et al. 2011a). Recently it has been demonstrated that *b_6_f* complex isolated from both spinach and *Mastigocladus* produce 20-30x more O_2_^•−^, on a per complex basis, than does the yeast cytochrome *bc*_1_ complex (Baniulis et al. 2013). Two potential O_2_^•−^ production sites were suggested, either the heme *b_p_* or PQ_*p*_^•−^ The reported E_0_’ of the heme *b_p_* is significantly more negative (−80 to −172 mV, (Hurt and Hauska 1983, 1982)) than those reported for yeast and mammalian cytochrome *bc_1_* heme *b_p_* (≈ −30 mV, (T’Sai and Palmer 1983; Wikstrom 1973)). This would make dioxygen reduction more feasible in the photosynthetic complex (Sarewicz et al. 2010). It was also hypothesized that dioxygen reduction by PQ_*p*_^•−^ might be facilitated by a longer residence time of the semiquinone at the PQ_*p*_-binding site (Baniulis et al. 2013). Other investigators have suggested that the Rieske iron-sulfur protein is involved in O_2_^•−^ production in both cytochrome *bc_1_* complexes (Genova et al. 2001) and the *b_6_f* complex (Sang et al. 2011a). In this regard it is interesting that Sang et al. (Sang et al. 2011a) reported that while no O_2_^•−^ was generated from complexes lacking the Rieske cluster, ^1^O2 was produced. These authors suggested that O_2_^•−^ was produced by, at least partially, a ^1^O_2_-dependent process (Sang et al. 2011a; Sang et al. 2011b).

The possible production of ^1^O_2_ by the *b_6_f* complex is intriguing. As noted above, the complex contains Chl *a*. The presence of chlorophyll prosthetic groups can be quite hazardous due to the possible production of ^1^O_2_ by intersystem crossing. Typically, chlorophylls are found in close proximity to carotenoids that can quench ^1^O_2_. The β-carotene in the *b_6_f* complex, however, is located ≥ 14 Å from the Chl *a* and too distant to serve as an efficient quencher (Dashdorj et al. 2005; Kim et al. 2005). It has been suggested that quenching of the ^1^O_2_ may be facilitated by a putative hydrophobic ROS channel which funnels ^1^O_2_ to the carotenoid (Kim et al. 2005). It should also be noted that aromatic residues in the vicinity of the Chl *a* significantly shorten the fluorescence lifetime (by about 20-fold), which would lower the yield of^3^ Chl, reducing the probability of ^1^O_2_ formation (Peterman et al. 1998; Dashdorj et al. 2005; Yan et al. 2008). Both Chl *a* and the β-carotene may also function in complex assembly (Cramer et al. 2009). Finally, it has been suggested that iron-sulfur proteins, in general, can serve as blue-light sensitizers for the production of ^1^O_2_ (Kim and Jung 1992). These authors have specifically suggested that the Rieske iron-sulfur cluster in the *b_6_f* complex is a major source of ^1^O_2_ in thylakoid membranes (Suh et al. 2000). This hypothesis is controversial, and strong evidence indicating that Chl *a* is the principal source of ^1^O_2_ has been presented (Sang et al. 2010).

It should also be noted that heme *c_n_* may directly bind dioxygen. This is suggested by the observation that NO, a dioxygen analogue, binds tightly to the heme (Twigg et al. 2009). The authors presented the possibility that heme *cn* could function as a plastoquinol oxidase. In this capacity an aberrant formation of O_2_^•−^ by a one-electron reduction of dioxygen (or H_2_O_2_ by a two-electron reduction) could hypothetically occur. It is also possible that the observed strong binding of NO, itself, is physiologically relevant as NO, often in cooperation with ROS, is involved in a wide variety of signal transduction pathways involving plant response to abiotic stress (Farnese et al. 2016; Mittler 2016).

No in-depth characterization of oxidative modification sites on the *b_6_f* complex has been performed and the relative importance and/or contribution of the different proposed ROS production sites (heme *b_p_*, PQ_*p*_, Fe_2_S_2_, Chl *a*) have not been evaluated. Galetskiy et al. (Galetskiy et al.) did report that eleven residues were ROS-modified within the complex but did not provide their locations. Importantly, these authors did not utilize a non-oxidizing denaturing PAGE system (see below) in their study. Consequently the possibility of protein oxidative modification artifacts due to electrophoretic conditions cannot be excluded.

In our study, we have used high-resolution tandem mass spectrometry to identify the location of oxidized residues within the cytochrome *b_e_f* complex isolated from field-grown spinach. These “natively” oxidized residues are the product of ROS production in the field environment where the plants may be exposed to a variety of abiotic stressors such as high light intensities, low or high temperatures, drought, etc. (You and Chan 2015; Choudhry et al. 2016).

It should be noted that earlier, we have used these methods to identify natively oxidized residues in spinach PS II (Frankel et al. 2012, 2013), results which have recently been confirmed and extended for the cyanobacterial photosystem (Weisz et al. 2017).

In the current study have mapped the natively oxidized residues identified in field-grown spinach cytochrome *b_e_f* complex onto the corresponding residues of the *Chlamydomonas b_6_f* complex structure (Stroebel et al. 2003). Our results indicate that numerous oxidized amino acid residues are located in the vicinity of the *p*-side cofactors heme *b_p_*, the Rieske iron-sulfur protein and the PQ_*p*_-binding site. None were observed in the vicinity of *n*-side cofactors. Additionally, oxidized residues were located adjacent to the Chl *a*. Our findings support the hypothesis that the *p*-side cofactors and Chl *a* are responsible for most of the ROS produced by the cytochrome *b_6_f* complex.

## Materials and Methods

The cytochrome *b_6_f* complex was isolated from market spinach essentially by the method previously described (Hurt and Hauska 1981). The *b_6_f* subunits were resolved on a 12.5-20% acrylamide gradient by LiDS-PAGE (Delepelaire and Chua 1979) either using the standard method (see Fig. 1B) or, for mass spectrometry, using a non-oxidizing gel system (Rabilloud et al. 1995). This was required, as standard PAGE is known to introduce numerous protein oxidation artifacts (Sun and Anderson 2004). In this system, after degassing, the gels are polymerized with riboflavin in the presence of diphenyliodonium chloride and toluensulfinate followed by exposure to UV light. The upper reservoir buffer contained thioglycolate.

**Figure 1.**
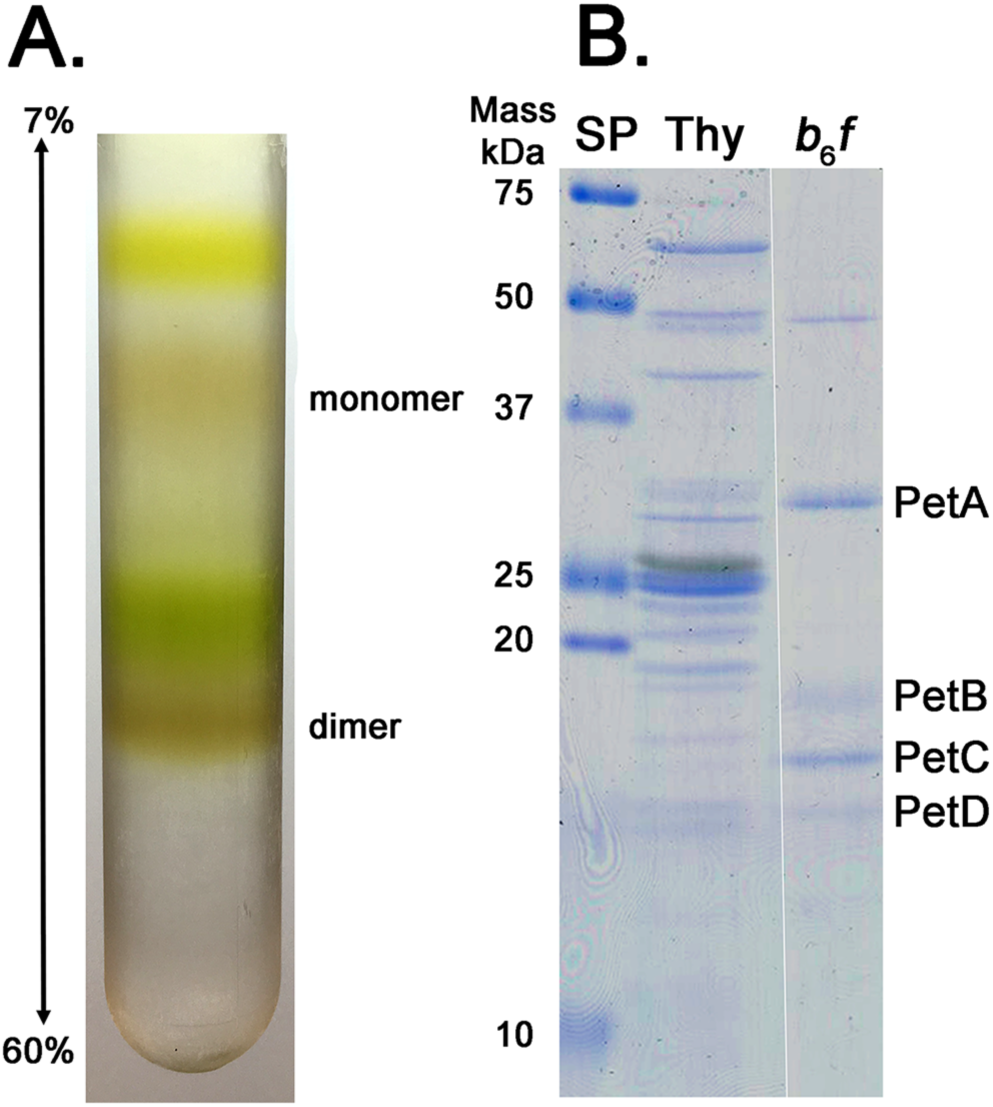
Purification of the Spinach Cytochrome *b_6_f* Complex. The complex was prepared essentially according to the methods of Hurt and Hauska (Hurt and Hauska 1981). A. Sucrose density gradient. Both monomer and dimer bands were observed; the dimer was used for all subsequent studies. B. LiDS-PAGE of thylakoids (Thy) and the *b_6_f* complex dimer. Subunits are labeled to the right, standard proteins to the left. The small subunits, PetG, PetL, PetM, and PetN which have apparent molecular masses in the 2-4 kDa region, are not resolved in this gel system.

Previously, we had demonstrated that PS II proteins resolved in this system exhibited much lower levels of artifactual protein oxidation than proteins resolved by standard PAGE (Frankel et al. 2012) confirming the earlier reports of Rabilloud et al. (Rabilloud et al. 1995) and Sun et al. (Sun and Anderson 2004), both of which examined other test proteins. For mass spectrometry, electrophoresis was terminated when the stacked proteins first entered the resolving gel. The gel was then stained with Coomassie blue, destained, and the thick protein band containing the stacked *b_6_f* subunits was excised. These were then processed using either trypsin, chymotrypsin or pepsin digestion following standard procedures. Three biological replicates were analyzed for each of the three proteases (chymotrypsin, pepsin and trypsin), and the union set of these replicates is presented. After protease digestion, the peptides were resolved by HPLC on a C:18 reversed phase column and ionized via electrospray into a Thermo Scientific Orbitrap Fusion Lumos mass spectrometer. The samples were analyzed in a data-dependent mode with one Orbitrap MS^1^ scan acquired simultaneously with up to ten linear ion trap MS^2^ scans.

Identification and analysis of the peptides containing oxidative modifications were performed using the MassMatrix Program (Xu and Freitas 2009). A library containing the sequences of the eight subunits of the spinach complex plus Ferredoxin-NADP+ oxidoreductase was searched, as was a decoy library which contained these same sequences but in reversed amino acid order. Twelve different types of oxidative modifications were included as possible post-translational modifications. For a positive identification of an oxidized residue, the peptide must exhibit a *p*-value of 10^−5^ or smaller; this value was selected prior to data collection. The identified oxidized amino acid residues were mapped onto the crystal structure of the *Chlamydomonas reinhardtii b_6_f* complex (PDB: 1Q90,(Stroebel et al. 2003)) using PYMOL (DeLano 2002).

## Results and Discussion

Isolation of the spinach *b_6_f* complex yielded results which were basically indistinguishable from previous reports (Hurt and Hauska 1981; Black et al. 1987; Zhang et al. 2001) for the isolation of the spinach complex (Fig. 1A). Four major polypeptides were identified: PetA, PetB, PetC and PetD (Fig. 1B). It should be noted that in standard LiDS-PAGE (Fig. 1B) the low molecular mass subunits (2-4 kDa) PetG, PetL, PetM, and PetN are not resolved. A fifth unidentified peptide with an apparent molecular mass of 48 kDa was also observed. This component, which is probably a contaminant, has been sporadically observed in other preparations of the complex (Hauska 2004). Tandem mass spectrometry analysis of the chymotryptic, peptic and tryptic peptides of the cytochrome *b_e_f* complex allowed the identifiication of 53 oxidatively modified residues present on the these subunits. The identity of these oxidized residues and the types of modifications observed are presented in Table 1. No oxidative modifications were observed on the small subunits of the complex (PetG, PetL, PetM and PetN). It should be emphasized that it is highly unlikely that all of the observed modifications would be present on every copy of the complex. Rather, this portfolio of detectable modifications is present within the full population of cytochrome *b_6_f* complexes present in our biological samples. It should be noted that for this study we isolated the complex from field-grown market spinach. Consequently, the exact growth conditions are unknown. Many studies examining the spinach cytochrome *b_6_f* complex have utilized comparable biological materials (Baymann et al. 2007; Szymańska et al. 2010; Stofleth 2012; Baniulis et al. 2013).

**Table 1.**
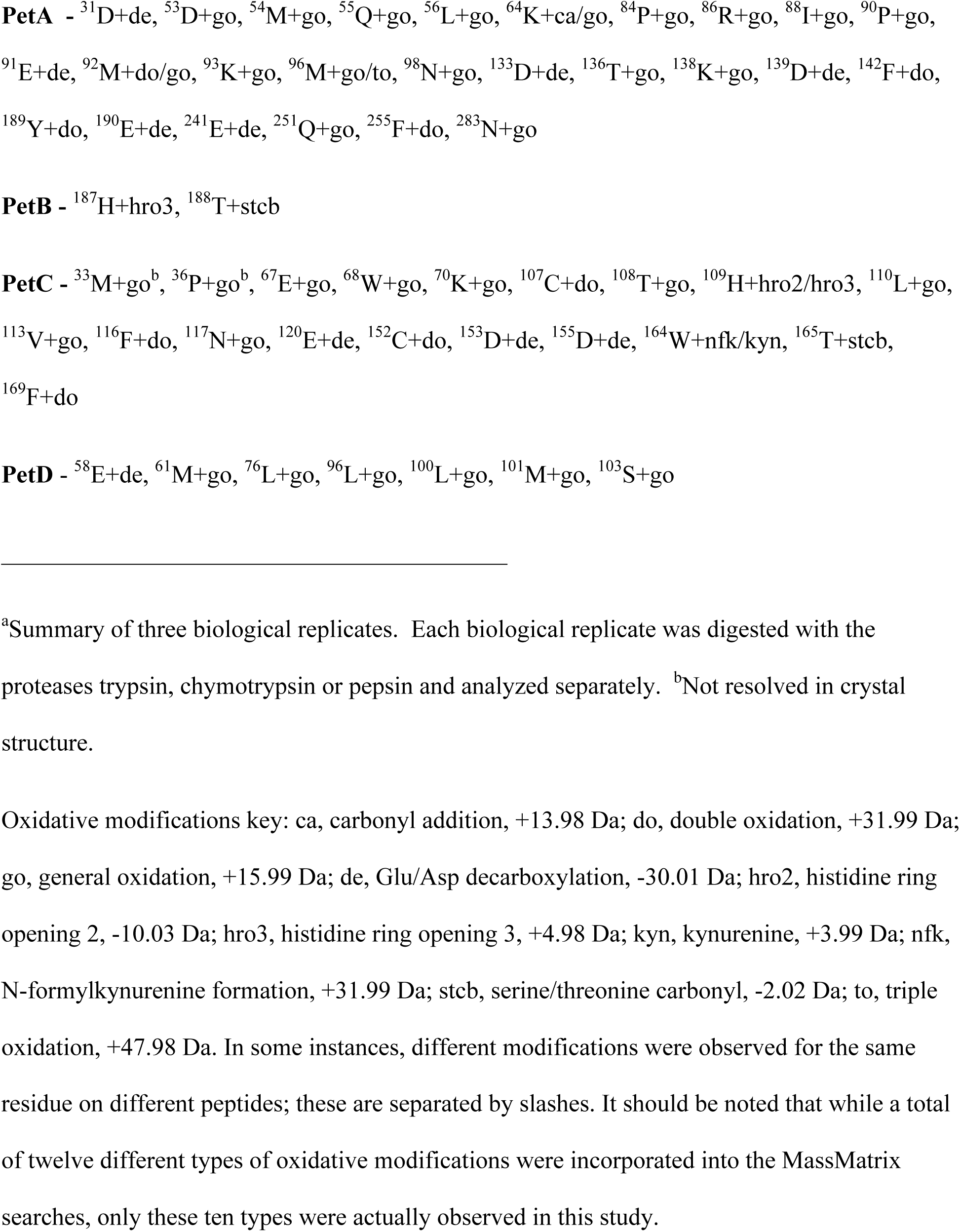
Natively Oxidized Residues in the Spinach *b_6_f* Complex^a^

Fig. 2 illustrates the quality of the data used for the identification of oxidized amino acid residues within the cytochrome *b_6_f* complex. In this figure, the tandem mass spectrometry data collected for the ^45^E-^56^L peptic peptide of PetA are illustrated. In Fig. 2A, the data from the unmodified peptide are shown, while in Fig. 2B, data from this peptide bearing oxidized ^54^M are shown. Both of these were observed in the same biological replicate. The observed mass accuracies for the parent peptic ions were −0.35 ppm and +0.34 ppm, respectively. The *p*-value for both of the illustrated peptides was 10^−55^ and are, consequently, among the lowest quality peptides used in this study (*p*-value range = 10^−50^ – 10^−11.1^). Even these peptides, however, clearly exhibited nearly complete y- and b-ion series allowing unequivocal identification of the oxidative mass modification. This result indicates that the use of *p* values ≤ 10^−5^ provided very high quality peptide identifications. Fig. S1 illustrates results for peptides exhibiting the median and lowest *p*-value peptides identified in this study ( *p*-values of 10^6^'^4^ and 10^−111^, respectively). One should note that all of the subunits of the cytochrome *b_6_f* complex are intrinsic membrane proteins. The analysis of such proteins by mass spectrometry is often difficult, with only relatively low sequence coverage being reported in standard “bottom-up” experiments (Souda et al. 2011; Kar et al. 2017; Weisz et al. 2017). In this study, however, we have obtained nearly complete coverage (>90%) for all of the major subunits of the complex using the enzymes trypsin, chymotrypsin and pepsin for proteolysis. This is illustrated in Fig. 3.

**Figure 2.**
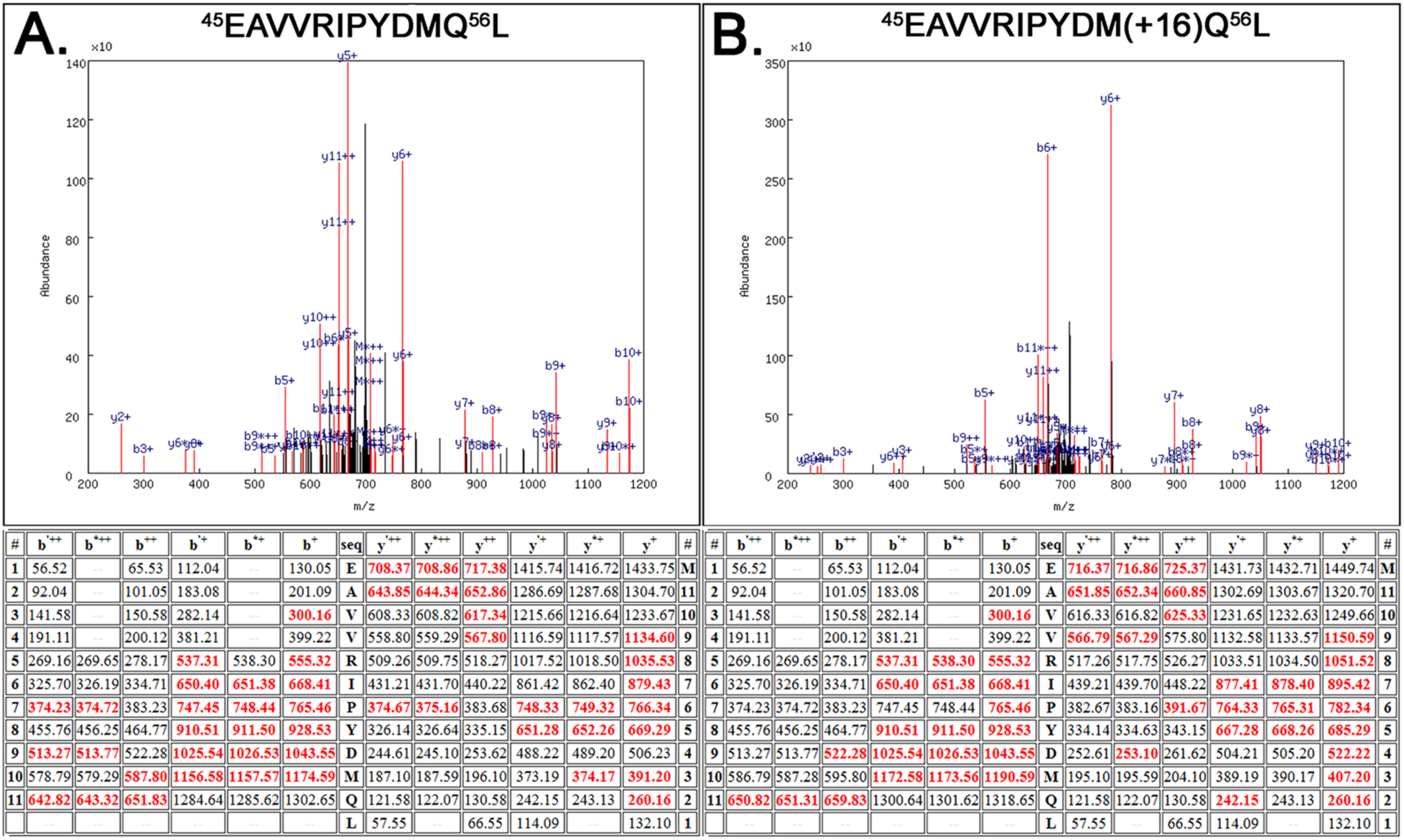
Quality of the Mass Spectrometry. Shown are the mass spectrometry results for the peptide PetA:^45^EAVVRIPYDMQ^56^L in both the unmodified (A) and modified (^54^M+16) forms (B). A. Top, spectrum of the CID dissociation of the unmodified peptide PetA:^45^EAVVRIPYDMQ^56^L. Various identified ions are labeled. Bottom, table of all predicted masses for the y- and b-ions generated from this peptide sequence. Ions identified in the CID spectrum (Top) are shown in red. The b’^++^, b’^+^ y’^++^ and y’^+^ ions are generated by the neutral loss of water while the b*^++^, b*^+^ y*^++^ and y*^+^ ions are generated from the loss of ammonia. B. Top, spectrum of the CID dissociation of the modified PetA:^45^EAVVRIPYDM(+16)Q^56^L. Various identified ions are labeled. Bottom, table of all predicted masses for the y- and b-ions generated from this peptide sequence. Ions identified in the CID spectrum are shown in red. The ions y3^+^-y9+ exhibit the +16 mass modification as does the b10^+^ ion when compared to the same ions in A. This verifies that ^54^M contains an oxidative modification. The *p*-values for both the unmodified and modified peptides were 10^-55^.

**Figure 3.**
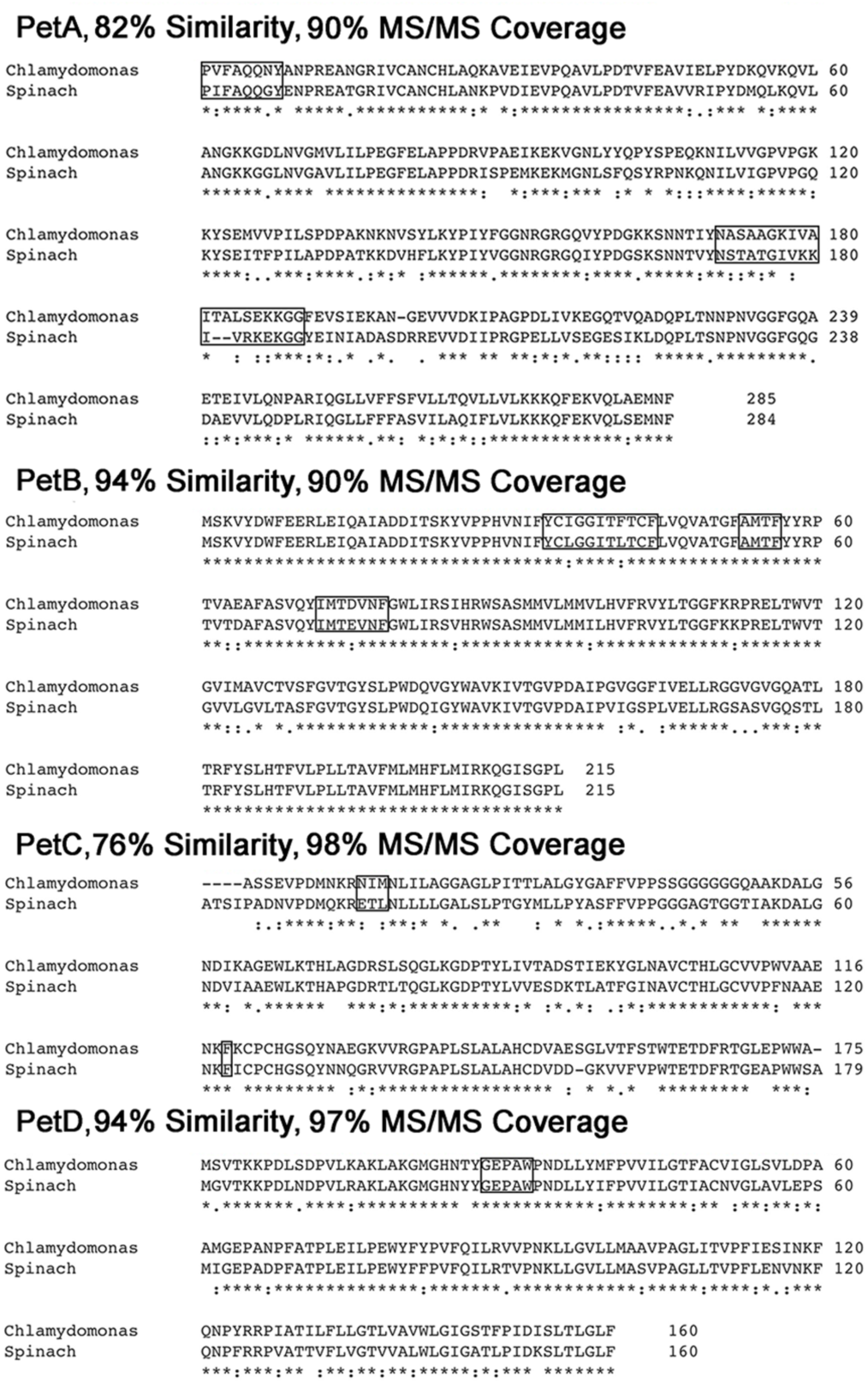
Sequence Alignments of Spinach and *Chlamydomonas* Cytochrome *b_6_f* Subunits and Mass Spectrometry Coverage. The subunits of the spinach and *Chlamydomonas* are very similar, which supports the use of the *Chlamydomonas* structure for these studies. Alignments were performed with CLUSTAL Omega (Sievers et al. 2011). Similarity scores were calculated using BLAST (Camacho et al. 2009). Combined mass spectrometry coverage of the *b_6_f* complex subunits, using trypsin, chymotrypsin, and pepsin coupled with Orbitrap analysis, was excellent (≥ 90%). Sequences which were not identified by mass spectrometry are boxed.

No crystal structure is currently available for the spinach cytochrome *b_6_f* complex. However, crystal structures are available for the thermophilic cyanobacteria *Mastigocladus laminosus* (Kurisu et al. 2003) and *Nostoc* sp. PCC7120 (Baniulis et al. 2009), as well as the mesophilic eukaryote *Chlamydomonas reinhardtii* (Stroebel et al. 2003). The sequence similarity between the spinach subunits and the *Chlamydomonas* subunits is high (Fig. 3), being 82% for PetA, 94% for PetB, 76% for PetC and 94% for PetD. This high degree of similarity allowed us to rationally map the oxidized amino acids that we observed in the spinach *b_6_f* complex onto the crystal structure of the *Chlamydomonas* protein complex. Indeed, 36 of the identified 53 modified residues (68%) were identical in both systems.

Fig. 4 presents an overview of the locations of the oxidized residues that we identified within the context of the cytochrome *b_6_f* complex dimer. The vast majority of the observed oxidized residues were located on the *p*-side of the complex. This does not appear to be the result of a sampling error since our mass coverage of the *n*-side residues was 96% (136/142 residues). Surface domains on the PetA and PetC subunits appear to be particularly susceptible to oxidative modification. This is not surprising since the surfaces of these components are exposed to the bulk solvent of the lumen. ROS produced by PS II due to manganese cluster damage (HO^·^ and, possibly, H_2_O_2_), ^1^O_2_ produced at P_680_*, the production of O_2_^•−^ and possibly other ROS species by the *b_6_f* complex itself, and possibly PS I, may all contribute to the oxidative modification of lumenally exposed domains. Additionally, while there are many ROS detoxification systems localized to the *n*-side of the thylakoid membrane (Tripathy and Oelmuller 2012; Das and Roychoudhury 2014), only a few lumenal components of putative *p*-side ROS detoxification systems have been reported (Levesque-Tremblay et al. 2009; Bermudez et al. 2012). Consequently, it is possible that ROS are not detoxified as efficiently in the thylakoid lumen as they are in the chloroplast stroma.

**Figure 4.**
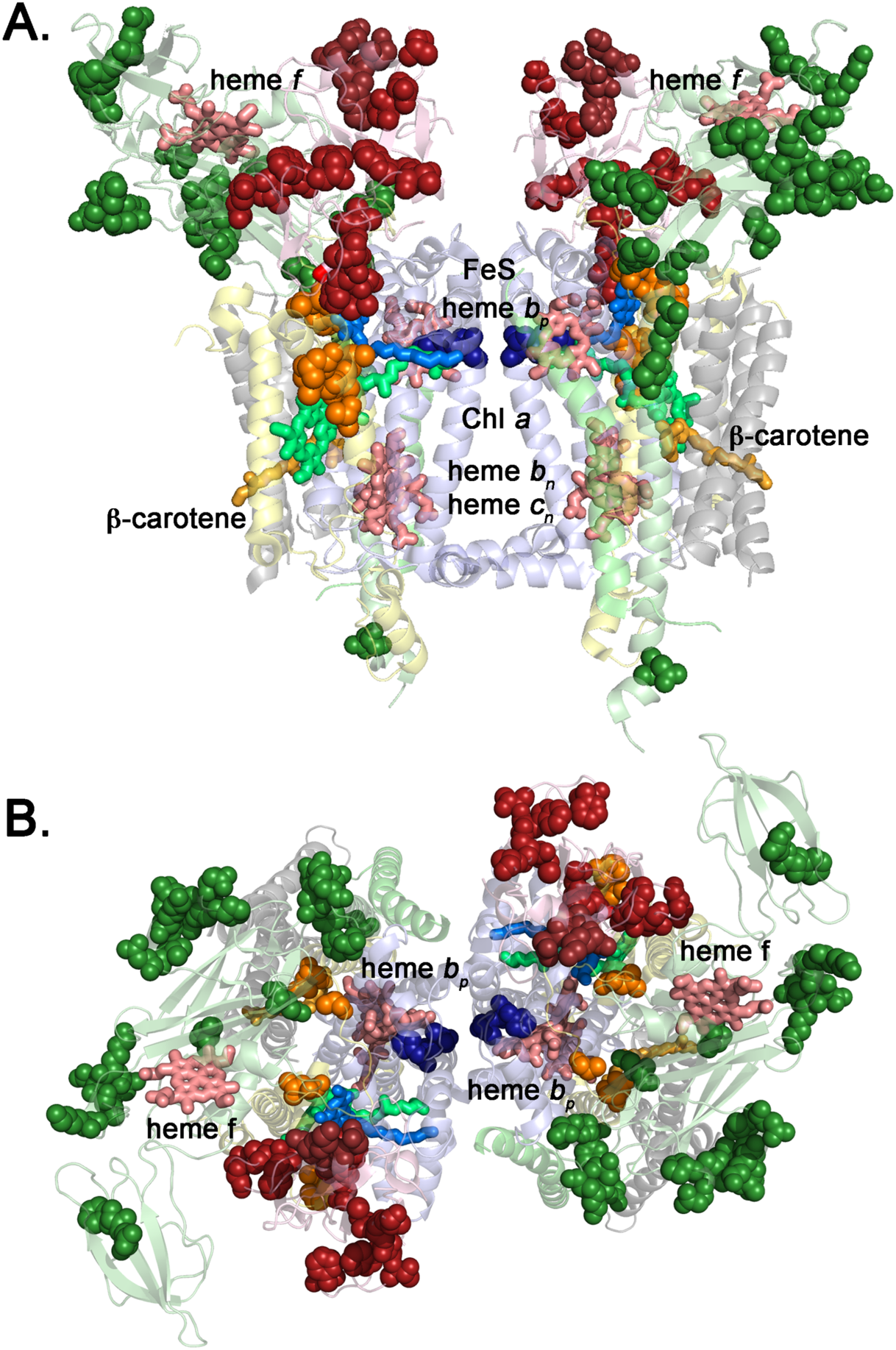
Overview of Natively Oxidized Amino Acid Residues in the Spinach *b_6_f* Complex. A. Side view of complex from within the plane of the membrane. B. Lumenal (*p*-side) view of the complex. The subunits are shown as follows: PetA (pale green), PetB (pale blue), PetC (pink), PetD (pale yellow) and the small subunits (grey). Oxidatively labeled residues are shown as spheres in darker shades and were mapped onto their corresponding locations on the *Chlamydomonas reinhardtii b_6_f* structure (Stroebel et al. 2003). Cofactors and TDS are shown in stick representation.

In addition to these surface-exposed oxidatively modified residues, a number of oxidized residues were observed which were buried or partially buried within the protein matrix, or present on the surface of the complex but buried within the lipid bilayer of the thylakoid membrane. Our working hypothesis is that amino acid residues that are in the vicinity of ROS production sites would be more prone to oxidative modification than residues that are more distant from these sites. In Fig. 5 we have examined the *p*-side cofactors which include: heme *f*, the Rieske iron-sulfur cluster, heme *b_p_* and the lumenal plastoquinol-binding site (PQ_*p*_) which, in this structure, is occupied by the *b_6_f* inhibitor TDS (tridecyl-stigmatellin). In Fig. 5, a 7.5 Å region surrounding each of the cofactors is shown with oxidized residues represented as spheres and labeled.

**Figure 5.**
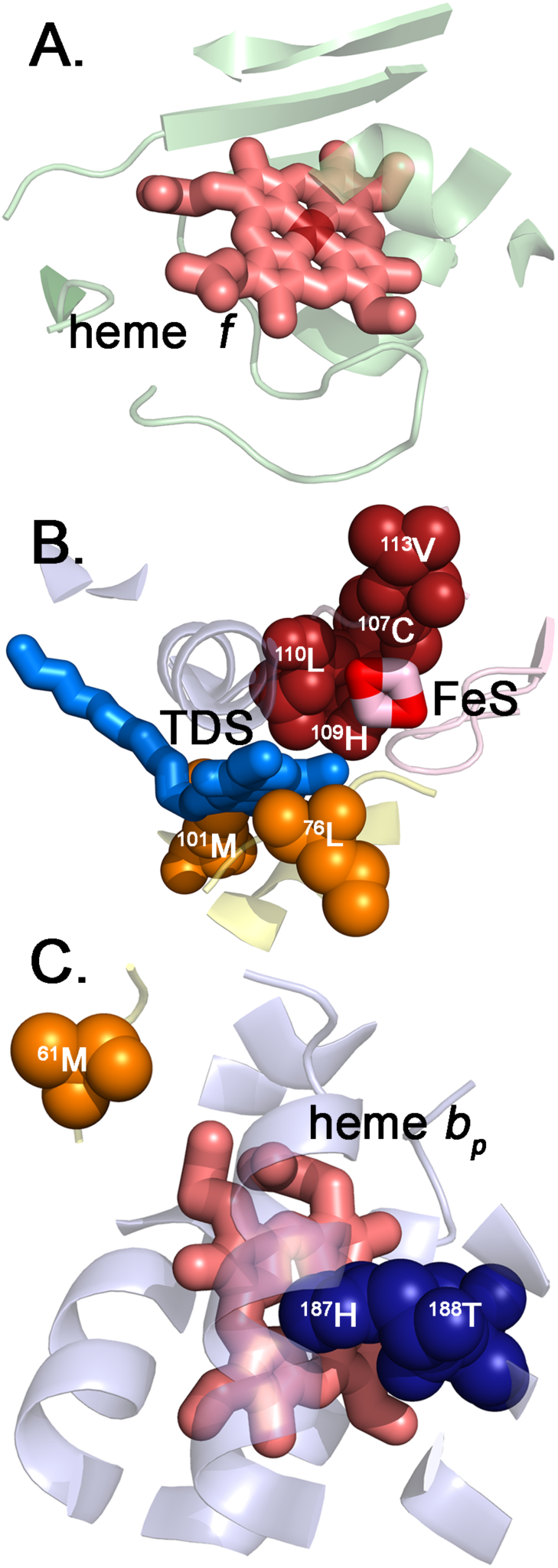
Details of the Oxidative Modifications Identified in the Vicinity of p-Side Cofactors. Shown is the protein structure located within 7.5 Å of the *p*-side cofactors cytochrome *f* (A), FeS and the PQ_*p*_-binding pocket, which is occupied by TDS (B), and cytochrome *b_p_* (C). Color coding of the *b_6_f* subunits is as shown in Fig. 4. Oxidatively modified residues are shown as spheres in darker shades and are labeled.

Fig. 5A demonstrates that even though a large number of oxidatively modified residues are located on the cytochrome*f* subunit, none of these are in close proximity to heme*f* This was expected since the production of ROS by this heme was not likely, given its high E_0_’ (+355 mV, (Alric et al. 2005)). Fig. 5B illustrates the location of oxidatively modified residues near the iron-sulfur cluster and the PQ_*p*_-binding site. The PetC residues ^107^C, ^108^T, ^109^H, ^110^L, and ^113^V were modified, as were ^76^L and ^101^M of PetD. The presence of seven oxidatively modified residues in close proximity to these cofactors strongly suggests that either the putative long-lived semiquinone occupying the PQ_*p*_-binding site and/or the iron-sulfur cluster is a source of ROS in the complex. The high E_0_’ (+320 mV) of the iron-sulfur cluster makes it unlikely that this site would be the source of O_2_^•−^. Additionally, the ability of the iron-sulfur cluster to act as a photosensitizer for ^1^O_2_ production is questionable (Sang et al. 2010). Nevertheless, we cannot rigorously exclude these possibilities at this time. Conversely, the ability of semiquinones to reduce O_2_ to O_2_^•−^ is well documented (Mubarakshina and Ivanov 2010). The production of O_2_^•−^ at the PQ_*p*_ site may be exacerbated by a hypothesized long residency time of the semiquinone (Baniulis et al. 2013). This long residency time might be due to the presence in the plastoquinol entrance/plastoquinone exit pathway of the phytyl tail of the Chl *a*, which might hinder quinol exchange.

In Fig. 5C, oxidized residues in the vicinity of heme *b_p_* are shown. Three residues, ^187^H and ^188^T of the PetB and ^61^M of PetD, were identified as being oxidatively modified. This observation raises the possibility that heme *bp* may also be a source of ROS, probably O_2_^•−^, as was previously hypothesized (Twigg et al. 2009; Sarewicz et al. 2010; Baniulis et al. 2013). Interestingly, ^187^H is a ligand to the heme iron. It is unclear what, if any, consequences this oxidative modification would have on the redox function of heme *b_p_.*

In Fig. 6 the immediate environment surrounding the hemes *b_n_* and *c_n_* are illustrated. No oxidatively modified residues were observed within 7.5 Å of the *b_n_* or *C_n_* hemes or the adjacent PQ_*n*_-binding pocket. It should be noted that in the*Mastigocladus* crystal structure (Hasan et al. 2013) the PQ_*n*_-binding pocket is occupied by TDS. This observation does not preclude the possibility that heme *c_n_* is associated with a putative plastoquinol oxidase activity (Twigg et al. 2009). It does suggest, however, that if an oxidase activity is present that it is efficient and not prone to the production of ROS in sufficient quantities to produce detectable oxidative modifications.

**Figure 6.**
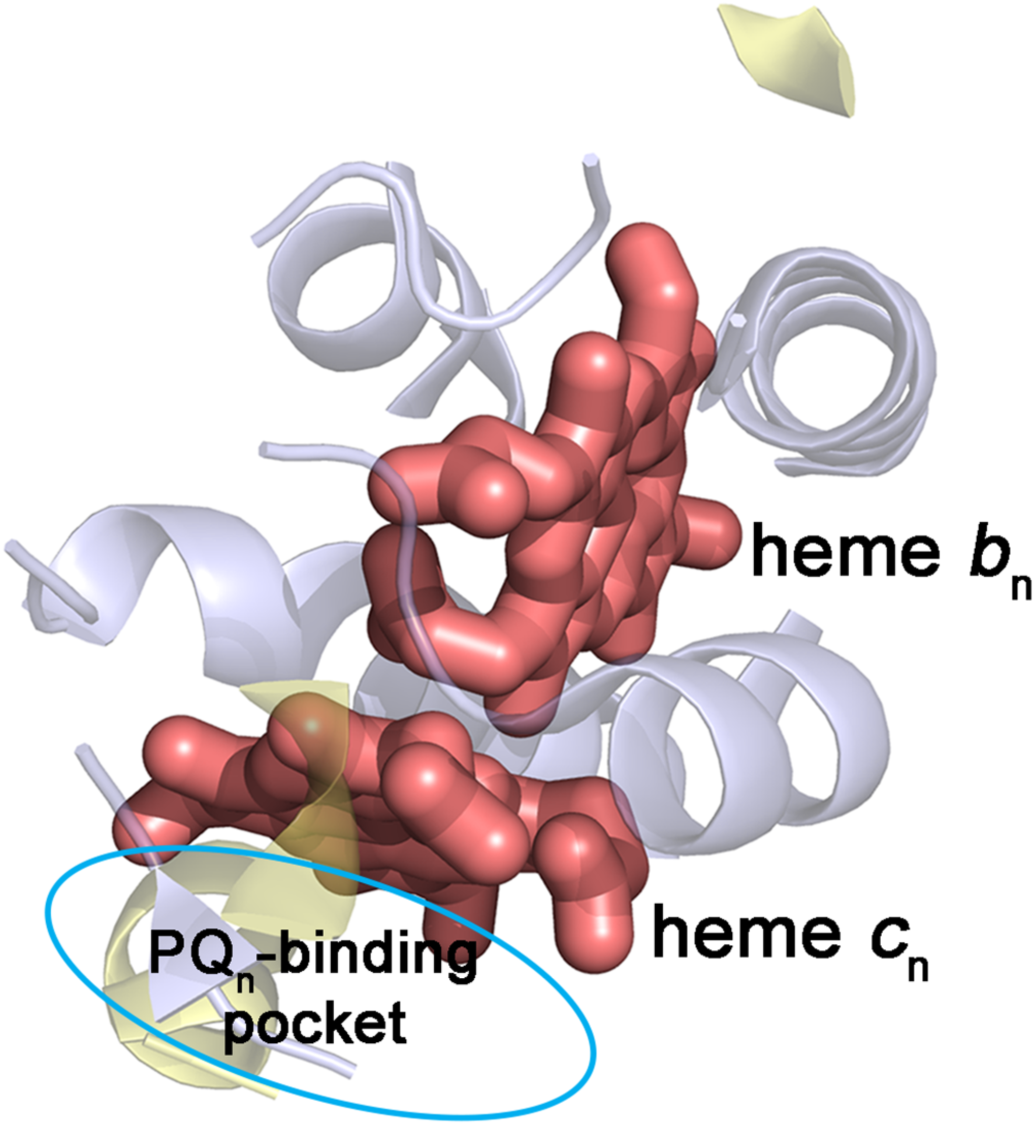
Details of the Oxidative Modifications Identified in the Vicinity of *n*-Side Cofactors. Shown is the protein structure located within 7.5 Å of the *n*-side cofactors cytochrome *b_n_* and cytochrome *c_n_*. The PQ_*n*_-binding pocket is indicated by a cyan ellipse. In the *Chlamydomonas* structure (Stroebel et al. 2003) this site is unoccupied while in the*Mastigocladus* structure it is occupied with TDS (Hasan et al. 2013).

In Fig. 7, oxidized residues in the vicinity of the Chl *a* and the β-carotene (Fig. 7A) are shown. Two residues adjacent to the Chl *a*, ^100^L and ^101^M of PetD (i.e. within 7.5 Å) are associated with the Chl a-binding pocket and, in the case of ^101^M, the PQp-binding site as well (see above). No oxidized residues were observed near the β-carotene. It had been hypothesized that a hydrophobic channel between the Chl *a* and the β-carotene exists which could funnel ^1^O_2_ from the Chl *a* to the β-carotene to facilitate quenching (Kim et al. 2005). This hypothetical channel would include residues PetB:^36^I,^95^L, ^96^M, ^98^I, ^99^L, and ^102^F, PetD:^133^F and several hydrophobic residues of PetG. None of these residues exhibited oxidative modification. It should be pointed out, however, that we do not have mass spectrometry coverage of PetB:^36^I and PetD:^133^F. Consequently, our results, while not precluding the presence of a channel, provide no evidence in support of this hypothesis. Interestingly, two other oxidized PetA residues were observed which are in contact with ^100^L and ^101^M; these residues, ^96^L and ^103^A (Fig. 7B), are also closely associated with the Chl a-binding pocket although more distant than 7.5 Å from the Chl *a* (12.1 and 8.8 Å, respectively). These results strongly suggest that ROS, probably ^1^O2, is produced at the Chl *a*, as has previously been hypothesized (Sang et al. 2010). It is possible that, since the Chl a-binding pocket is exposed at the surface of the complex but buried in the lipid bilayer, that ^1^O_2_ is released directly from the Chl a-binding pocket to the lipid bilayer (Fig. 7B).

**Figure 7.**
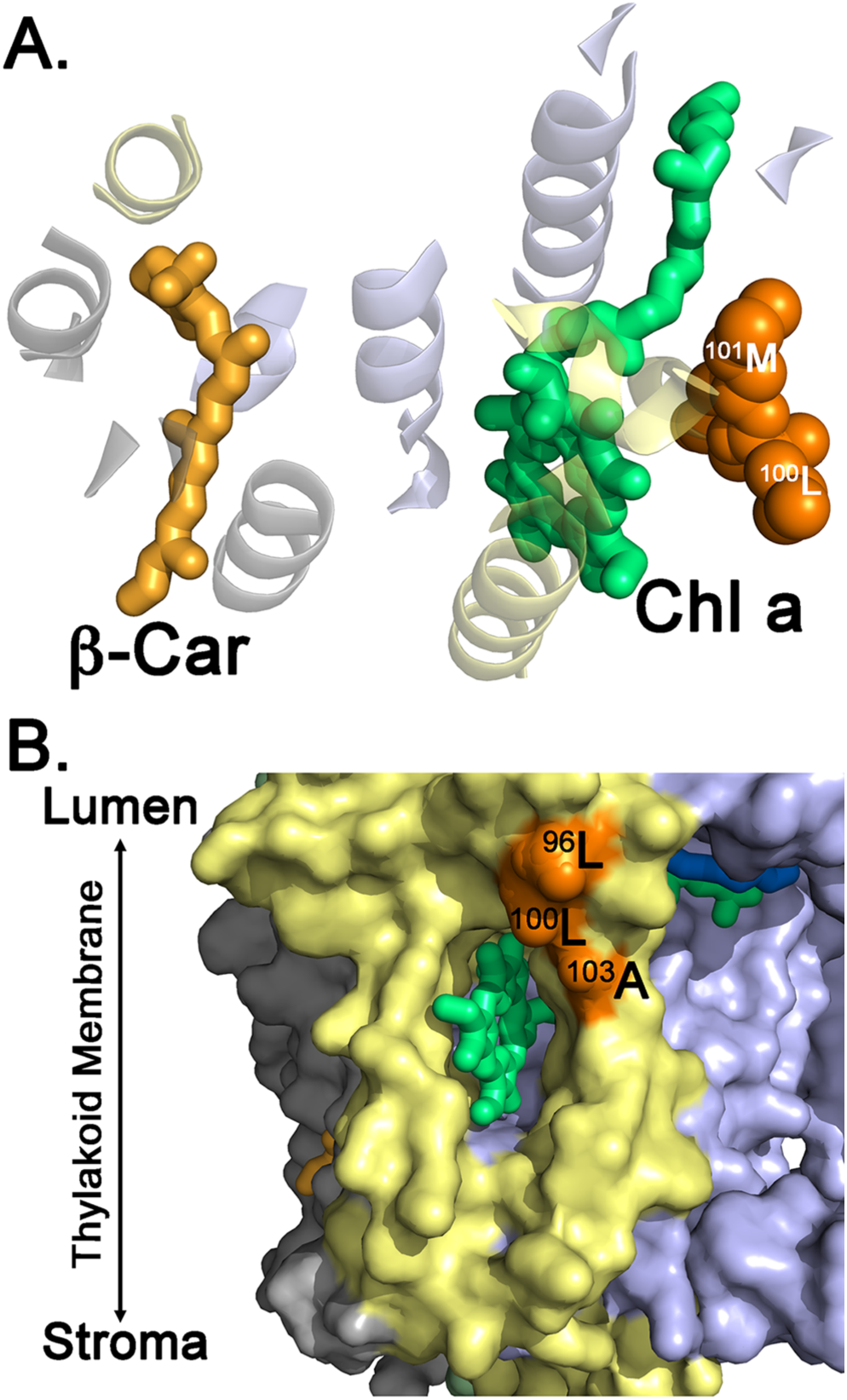
Details of the Oxidative Modifications Identified in the Vicinity of Chl *a* and the β-Carotene. A. Shown is the protein structure located within 7.5 Å of the Chl *a* and the β-carotene. Color coding of the *b_6_f* subunits is as shown in Fig. 4. Oxidatively modified residues are shown as spheres in darker shades and are labeled. Note that no oxidative modifications were observed for the intervening hydrophobic residues located between the Chl *a* and the β-Carotene. B. Surface of the *b_6_f* complex in the vicinity of the Chl *a*-binding pocket. Color coding of the *b_6_f* subunits is as shown in Fig. 4. Oxidatively modified residues are shown as spheres in darker shades and are labeled.

## Conclusions

In this communication, we have identified numerous oxidized residues in the vicinity of the p-side cofactors heme *b_p_*, the Rieske iron sulfur cluster, the PQ_*p*_-binding domain and adjacent to the Chl *a*-binding pocket. The locations of these modified residues are consistent with our hypothesis that residues in the vicinity of ROS production sites would be prone to oxidative modification. We have not, at this time, determined the type of ROS leading to these observed modifications, the relative importance of the various possible sites in ROS production nor the time course for the appearance of oxidative modifications. These important questions are the subject of future studies.

## Acknowledgements

This work was supported by the United States Department of Energy, Office of Basic Energy Sciences grant DE-FG02-09ER20310 to TMB and LKF.

## References

1. Kurisu G, Zhang H, Smith JL, Cramer WA (2003) Structure of the cytochrome b6 *f* complex of oxygenic photosynthesis: tuning the cavity. Science 302: 1009–1014.

2. Baniulis D, Yamashita E, Whitelegge JP, Zatsman AI, Hendrich MP, et al. (2009) Structure-function, stability, and chemical modification of the cyanobacterial cytochrome *b_6_f* Complex from *Nostoc* sp. PCC 7120. J Biol Chem 284: 9861–9869.

3. Stroebel D, Choquet Y, Popot JL, Picot D (2003) An atypical haem in the cytochrome *b_6_f* complex. Nature 426: 413–418.

4. Crofts AR, Shinkarev VP, Kolling DRJ, Hong S (2003) The modified Q-cycle explains the apparent mismatch between the kinetics of reduction of cytochromes *c_I_* and *b_H_* in the *bci* complex. J Biol Chem 278: 36191–36201.

5. Zhang H, Whitelegge JP, Cramer WA (2001) Ferredoxin NADP+ oxidoreductase is a subunit of the chloroplast cytochrome *b_6_f* complex. J Biol Chem 276: 38159–38165.

6. Yan J, Dashdorj N, Baniulis D, Yamashita E, Savikhin S, et al. (2008) On the structural role of the aromatic residue environment of the chlorophyll *a* in the cytochrome *b_6_f* complex. Biochemistry 47: 3654–3661.

7. Das K, Roychoudhury A (2014) Reactive oxygen species (ROS) and response of antioxidants as ROS-scavengers during environmental stress in plants. Front Environ Sci 2: 53.

8. Long SP, Humphries S, Falkowski PG (1994) Photoinhibition of photosynthesis in nature. Ann Rev Plant Mol Biol 45: 633–662.

9. Mittler R (2016) ROS are good. Trends in Plant Science 7, 405–410.

10. Kim CS, Jung J (1992) Iron-sulfur centers as endogenous blue light sensitizers in cells: a study with an artificial non-heme iron protein. Photochem Photobiol 56: 63–68.

11. Pospíšil P (2009) Production of reactive oxygen species by Photosystem II. Biochim Biophys Acta 1787: 1151–1160.

12. Pospíšil P (2016) Production of reactive oxygen species by Photosystem II as a response to light and temperature stress. Front Plant Sci 7: 1950.

13. Kale R, Hebert AE, Frankel LK, Sallans L, Bricker TM, et al. (2017) Amino acid oxidation of the D1 and D2 proteins by oxygen radicals during photoinhibition of Photosystem II. Proc Natl Acad Sci (USA) 114: 2988–2993.

14. Lanciano P, Khalfaoui-Hassani B, Selamoglu N, Ghelli A, Rugolo M, et al. (2013) Molecular mechanisms of superoxide production by complex III: A bacterial versus human mitochondrial comparative case study. Biochim Biophys Acta 1827: 1332–1339.

15. Sang M, Qin X-C, Wang W-D, Xie J, Chen X-B, et al. (2011) High-light-induced superoxide anion radical formation in cytochrome *b_6_f* complex from spinach as detected by EPR spectroscopy. Photosynthetica 49: 48–54.

16. Baniulis D, Hasan SS, Stofleth JT, Cramer WA (2013) Mechanism of enhanced superoxide production in the cytochrome *b_6_f* complex of oxygenic photosynthesis. Biochemistry 52: 8975–8983.

17. Sarewicz M, Borek A, Cieluch E, Swierczek M, Osyczka A (2010) Discrimination between two possible reaction sequences that create potential risk of generation of deleterious radicals by cytochrome *bc*_1_. Implications for the mechanism of superoxide production. Biochim Biophys Acta 1797: 1820–1827.

18. Genova ML, Ventura B, Giuliano G, Bovina C, Formiggini G, et al. (2001) The site of production of superoxide radical in mitochondrial Complex I is not a bound ubisemiquinone but presumably iron-sulfur cluster N2. FEBS Lett 505: 364–368.

19. Sang M, Xie J, Qin X-C, Wang W-D, Chen X-B, et al. (2011) High-light induced superoxide radical formation in cytochrome *b_6_f* complex from *Bryopsis corticulans* as detected by EPR spectroscopy. Photochem Photobiol 102: 177–181.

20. Kim H, Dashdorj N, Zhang H, Yan J, Cramer WA, et al. (2005) An anomalous distance dependence of intra-protein chlorophyll-carotenoid triplet energy transfer. Biophys J 89: PL28–L30.

21. Peterman EJG, Wenk S-O, Pullerits T, Pålsson L-O, van Grondelle R, et al. (1998) Fluorescence and absorption spectroscopy of the weakly fluorescent chlorophyll *a* in cytochrome *b_6_f* of *Synechocystis* PCC6803. Biophys J 75: 389–398.

22. Dashdorj N, Zhang H, Kim H, Yan J, Cramer WA, et al. (2005) The single chlorophyll *a* molecule in the cytochrome *b_6_f* complex: unusual optical properties protect the complex against singlet oxygen. Biophys J 88: 4178–4187.

23. Cramer WA, Savikhin S, Yan J, Yamashita E (2009) The enigmatic chlorophyll *a* molecule in the cytochrome *b_6_f* complex. In: Rebeiz C, Bohnert H, Benning C, Daniell H, Hoober K et al., editors. The Chloroplast System: Biochemistry and Molecular Biology. Dordrecht: Springer. pp. 89–92.

24. Suh H-J, Kim CS, Jung J (2000) Cytochrome *b_6_/f* complex as an indigenous photodynamic generator of singlet oxygen in thylakoid membranes. Photochem Photobiol 71: 103–109.

25. Sang M, Ma F, Chen X-B, Wang KB, Qin X-C, et al. (2010) High-light induced singlet oxygen formation in cytochrome *b_6_f* complex from *Bryopsis corticulans* as detected by EPR spectroscopy. Biophys Chem 146: 7–12.

26. Twigg AI, Baniulis D, Cramer WA, Hendrich MP (2009) EPR detection of an O_2_ surrogate bound to heme *c_n_* of the cytochrome *b_6_f* complex. J Amer Chem Soc 131: 12536–12537.

27. Farnese FS, Menezes-Silva PE, Gusman GS, Oliveira JA (2016) When Bad Guys Become Good Ones: The Key Role of Reactive Oxygen Species and Nitric Oxide in the Plant Responses to Abiotic Stress. Front Plant Sci 7: 471

28. Galetskiy D, Lohscheider JN, Kononikhin AS, Popov IA, Nikolaev EN, et al. (2011) Mass spectrometric characterization of photooxidative protein modifications in *Arabidopsis thaliana* thylakoid membranes. Rapid Commun Mass Spectrom 25: 184–190.

29. Hurt E, Hauska G (1981) A cytochrome *f/b_6_* complex of five polypeptides with plastoquinol-plastocyanin-oxidoreductase activity from spinach chloroplasts. Eur J Biochem 117: 591–599.

30. Delepelaire P, Chua NH (1979) Lithium dodecyl sulfate/polyacrylamide gel electrophoresis of thylakoid membranes at 4 degrees C: Characterizations of two additional chlorophyll a-protein complexes. Proc Natl Acad Sci U S A 76: 111–115.

31. Rabilloud T, Vincon M, Garin J (1995) Micropreparative one- and two-dimensional electrophoresis: Improvement with new photopolymerization systems. Electrophoresis 16: 1414–1422.

32. Sun G, Anderson VE (2004) Prevention of artifactual protein oxidation generated during sodium dodecyl sulfate-gel electrophoresis. Electrophoresis 25: 959–965.

33. Frankel LK, Sallans L, Limbach PA, Bricker TM (2012) Identification of oxidized amino acid residues in the vicinity of the Mn_4_CaO_5_ cluster of Photosystem II: Implications for the identification of oxygen channels within the photosystem. Biochemistry 51: 6371–6377.

34. Xu H, Freitas MA (2009) MassMatrix: A database search program for rapid characterization of proteins and peptides from tandem mass spectrometry data. Proteomics 9: 1548–1555.

35. DeLano WL (2002) The PyMOL molecular graphics system. Software.

36. Black MT, Widger WR, Cramer WA (1987) Large-scale purification of active cytochrome *b_6_/f* complex from spinach chloroplasts. Arch Biochem Biophys 252: 655–661.

37. Hauska G (2004) The isolation of a functional cytochrome *b_6_f* complex: from lucky encounter to rewarding experiences. Photosyn Res 80: 277–291.

38. Baymann F, Giusti F, Picot D, Nitschke W (2007) The *c_I_/b_H_* moiety in the *b_6_f* complex studied by EPR: A pair of strongly interacting hemes. Proc Natl Acad Sci (USA) 104: 519–524.

39. Szymańska R, Dluzewska J, Slesak I, Kruk J (2010) Ferredoxin:NADP+oxidoreductase bound to cytochrome *b_6_f* complex is active in plastoquinone reduction: Implications for cyclic electron transport. Physiol Plant 141: 289–298.

40. Stofleth JT (2012) Understanding free radicals: Isolating active thylakoid membranes and purifying the cytochrome *b_6_f* complex for superoxide generation studies. J Pur Undergrad Res 2: 64–69.

41. Tripathy BC, Oelmuller R (2012) Reactive oxygen species generation and signaling in plants. Plant Signal Behav 7: 1621–1633.

42. Levesque-Tremblay G, Havaux M, Ouellet F (2009) The chloroplastic lipocalin AtCHL prevents lipid peroxidation and protects Arabidopsis against oxidative stress. Plant J 60: 691–702.

43. Bermudez MA, Galmes J, Moreno I, Mullineaux PM, Gotor C, et al. (2012) Photosynthetic adaptation to length of day is dependent on S-sulfocysteine synthase activity in the thylakoid lumen. Plant Physiol 160: 274–288.

44. Alric J, Pierre Y, Picot D, Lavergne J, Rappaport F (2005) Spectral and redox characterization of the heme *ci* of the cytochrome *b_6_f* complex. Proc Natl Acad Sci (USA) 102: 15860–15865.

45. Mubarakshina MM, Ivanov BN (2010) The production and scavenging of reactive oxygen species in the plastoquinone pool of chloroplast thylakoid membranes. Physiol Plant 140: 103–110.

46. Hasan SS, Yamashita E, Baniulis S, Cramer WA (2013) Quinone-dependent proton transfer pathways in the photosynthetic cytochrome *b_6_/f* complex. Proc Natl Acad Sci (USA) 110: 4297–4302.

47. Sievers F, Wilm A, Dineen DG, Gibson TJ, Karplus K, et al. (2011) Fast, scalable generation of high-quality protein multiple sequence alignments using Clustal Omega. Mol Sys Biol 7: 539.

48. Camacho C, Coulouris G, Avagyan V, Ma N, Papadopoulos J, et al. (2009) BLAST+: architecture and applications. BMC Bioinform 10: 421–430.

